# Toxin-based screening of C-terminal tags in *Escherichia coli* reveals the exceptional potency of ssrA-like degrons

**DOI:** 10.1101/2024.01.29.576913

**Authors:** Patrick C. Beardslee, Karl R. Schmitz

## Abstract

All bacteria possess ATP-dependent proteases that destroy cytosolic proteins. These enzymes help cells mitigate proteotoxic stress, adapt to changing nutrient availability, regulate virulence phenotypes, and transition to pathogenic lifestyles. Moreover, ATP-dependent proteases have emerged as promising antibacterial and antivirulence targets in a variety of pathogens. The physiological roles of these proteases are largely defined by the complement of proteins that they degrade. Substrates are typically recognized in a highly selective manner, often via short unstructured sequences termed degrons. While a few degrons have been identified and rigorously characterized, we lack a systematic understanding of how proteases select valid degrons from the vast complexity of protein sequence space. Here, we describe a novel high-throughput screening approach in *Escherichia coli* that couples proteolysis of a protein toxin to cell survival. We used this method to screen a combinatorial library of C-terminal pentapeptide sequences for functionality as proteolytic degrons in wild type *E. coli*, and in strains lacking components of the ClpXP and ClpAP proteases. By examining the competitive enrichment of sequences over time, we found that about one percent of pentapeptide tags lead to toxin proteolysis. Interestingly, the most enriched degrons were ClpXP-dependent and highly similar to the ssrA tag, one of the most extensively characterized degrons in bacteria. Among ssrA-like sequences, we observed that specific upstream residues correlate with successful recognition. The lack of diversity among strongly enriched sequences suggests that ssrA-like tags comprise a uniquely potent class of short C-terminal degron in *E. coli*. Efficient proteolysis of substrates lacking such degrons likely requires adaptors or multivalent interactions. These findings broaden our understanding of the constraints that shape the bacterial proteolytic landscape. Our screening approach may be broadly applicable to probing aspects of proteolytic substrate selection in other bacterial systems.

## INTRODUCTION

Targeted degradation of cytosolic proteins by ATP-dependent proteases is critical for maintaining cell health throughout all domains of life [1]. Bacteria possess multiple ATP-dependent proteases that maintain protein quality control, regulate discrete pathways, and mitigate proteotoxic stress by destroying damaged or unneeded proteins [2]. In many pathogenic bacteria, these enzymes regulate virulence or are strictly essential for viability, and have thus emerged as promising drug targets [3, 4]. ATP-dependent proteolysis is notably decentralized in bacteria. *E. coli,* for example, possesses five structurally and functionally similar proteases: ClpXP, ClpAP, HslUV, Lon, and FtsH. All are large oligomeric complexes that consist of a ring-shaped ATP-dependent unfoldase that collaborates with a self-compartmentalized peptidase [5]. The active sites of the barrel-shaped peptidase are sequestered inside a solvent-filled chamber, thereby avoiding unregulated proteolysis of native proteins. Access to the inside of the peptidase is controlled by the unfoldase, which selects protein substrates, mechanochemically unfolds them, and translocates denatured polypeptides through its axial pore and into the peptidase for degradation [6]. Substrate recognition is therefore a crucial facet of proteolytic regulation, and largely defines the biological roles of proteases within the cell [7].

Bacterial ATP-dependent proteases have evolved a variety of mechanisms to selectively recognize substrates. For example, in *Caulobacter crescentus* a hierarchy of adaptors and anti-adaptors control substrate delivery to ClpXP during cell cycle progression [8–10]. In some firmicutes and actinobacteria, post-translational arginine phosphorylation marks proteins for destruction by ClpCP [11–13]. Additionally, many actinobacteria post-translationally modify proteins with prokaryotic ubiquitin-like protein (PUP), which is recognized as a degradation signal by the Mpa•20S proteasome [14, 15].

Whereas these mechanisms require the participation of auxiliary components, bacterial ATP-dependent proteases can also directly recognize some substrates via short unstructured sequence elements, termed degrons (**Fig. 1A**) [5]. Degrons have been identified at the N-termini, C-termini, and internal positions of substrates [7]. Some are constitutively exposed and program rapid proteolytic turnover [16–19], while others are conditionally revealed by a cleavage event or conformational change [20, 21]. One of the most well-studied degrons is the ssrA tag, a ∼10 amino acid sequence that is co-translationally appended to stalled polypeptides during tmRNA-mediated ribosomal rescue [22, 23]. Proteins bearing the ssrA degron are robustly proteolyzed by ClpXP, and to a lesser extent by other cellular proteases [24–27]. The *E. coli* ssrA tag (AANDENYALAA) has been extensively characterized and has serves as a versatile tool for studying the biochemical and mechanochemical properties of ClpXP and other proteases [28–32]. Furthermore, it has seen broad use in synthetic biology as a component in engineered protein degradation pathways [33–35].

**Figure 1.**
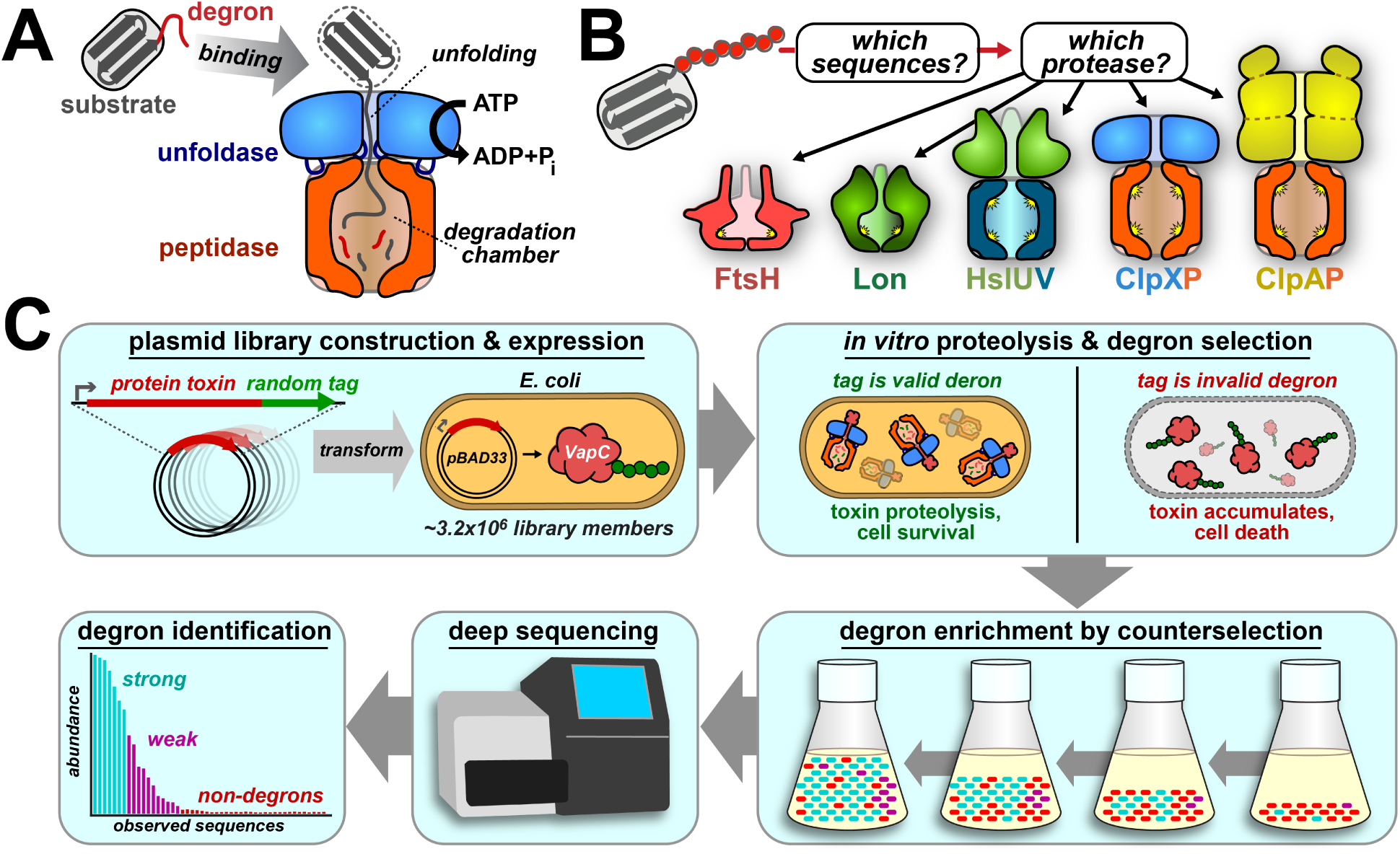
Coupling toxin proteolysis with cell survival. **A)** ATP-dependent proteases can initiate proteolysis by recognizing substrate degrons. **B)** Sequence-based recognition of substrates is poorly understood in bacteria. *E. coli*’s five ATP-dependent proteases are depicted. **C)** Overview of the DEtox screening approach, which couples degron-directed toxin proteolysis to degron enrichment in liquid culture.

While most ATP-dependent proteases have the ability to recognize degrons, relatively few specific degrons have been described. Several prior efforts have sought to systematically uncover substrates and degrons within individual bacterial species. These include approaches that trap substrates within inactivated peptidases or unfoldases, which have been carried out in *E. coli, Staphylococcus aureus, Caulobacter crescentus* and *Mycolicibacterium smegmatis* [36–41]. In *E. coli,* this method uncovered a handful of enriched N- and C-terminal motifs recognized by ClpXP or ClpAP. However, efforts to identify degrons within an endogenous proteome are inherently restricted to a pool of several thousand proteins, and thus sample only a tiny subset of the full sequence space encoded by even a short peptide library. More recently, a combinatorial approach was used to systematically screen all possible C-terminal dipeptides on a proteolytic reporter in *Mycoplasma pneumoniae*, revealing that that hydrophobic residues render proteins more susceptible to proteolysis [42]. Despite these and other findings, we currently lack a systematic understanding of the sequence-based rules governing selection of longer degrons. It remains an open question how many degron classes exist, how they vary in strength, and the extent to which they contribute to overall protein turnover [7] (**Fig. 1B**).

Here, we report a novel high-throughput selection-based screening platform for identifying degrons in bacteria that couples proteolysis of a toxin to cell survival. In our approach, which we term Degron Enrichment by Toxin (DEtox), *E. coli* are transformed with a plasmid library encoding a small protein toxin that bears a randomized C-terminal pentapeptide. Expression of the toxin places selective pressure on cells. Non-degron tags permit toxin accumulation, causing growth arrest. Tags that function as degrons promote toxin proteolysis, allowing cell proliferation. Cells expressing valid degrons are thus competitively enriched over time, and enrichment patterns can be read by next-generation sequencing (NGS) **(Fig. 1C**). We implemented this approach in wild-type *E. coli* and in several proteolytic deletion strains. In the resulting dataset, the majority of highly enriched sequences bore clear similarity to the ssrA tag, and strong enrichment only occurred when elements of the ClpXP protease were present. Surprisingly, we did not identify any non-ssrA-like sequences of similar strength. Our results indicate that ssrA-like sequences comprise a uniquely potent degron class, at least within the parameters of our screen, and that ClpXP plays the most significant role in recognizing short C-terminal degrons in *E. coli*. Moreover, our work suggests that short non-ssrA degrons are recognized weakly, reinforcing the importance of adaptors and other indirect recognition mechanisms in shaping proteolytic landscapes.

## MATERIALS & METHODS

### Strains & plasmids

All cell-based experiments and molecular cloning were carried out in *E. coli* strain MC1061 (Lucigen). Protein overexpression for purification was carried out in a derivative of *E. coli* strain ER2566 (NEB) harboring a deletion of *clpP*. Proteolytic deletion strains of MC1061 and ER2566 were generated by λ-Red recombineering [43]. Variants of VapC (Uniprot ID: E0J1H5) for *in vivo* expression were PCR-amplified from *E. coli* strain W [44] (ATCC) gDNA and cloned into pBAD33 [45] using BsaI endonuclease (NEB) and T4 DNA ligase (NEB). GFP variants were cloned downstream of a 7xHis-SUMO tag in a modified pET22b vector (EMD Millipore). Tag variants were generated by ligating synthetic dsDNA oligos (IDT) encoding each respective tag downstream of GFP bewteen BamHI and HindIII sites (NEB). In order to generate the GFP-VapC^YALAA^ fusion, VapC was PCR-amplified and cloned into the plasmid hosting GFP^YALAA^ by Gibson Assembly [46].

### Proteins

*^Eco^*ClpX^ΔN^ and *^Eco^*ClpP were expressed and purified as previously described [47]. For expression of GFP constructs, cultures were grown to OD_600_ ≈ 0.75 in 1.5xYT media (Genesee) at 37 °C, and induced with 500 µM IPTG for 4 h at 30 °C. Cells were harvested by centrifugation at 4000×g for 30 min at 4°C, and resuspended in 25 mL of Lysis Buffer (25 mM HEPES, 300 mM NaCl, 10 mM imidazole, 10% glycerol, pH 7.5). Cells were lysed by sonication and lysates were clarified by centrifuging for 1 h at 15,000×g. Proteins were purified by IMAC chromatography (Ni-NTA agarose; MCLabs), and size-exclusion chromatography (Superdex 200; Cytiva). SUMO tags were removed following Ni-NTA chromatography by incubating the pooled, concentrated elution fractions with the Ulp1 protease at a 1:50 ratio of Ulp1 to target protein for 2 hours at 30 °C [48, 49]. Cleaved proteins were exchanged into PD buffer (25 mM HEPES, 100 mM KCl, 10% glycerol, 5 mM MgCl_2_, 0.1 mM EDTA) over a Sephadex G-25 column (Cytiva) and separated from Ulp1 and the cleaved 7xHis-SUMO tag by additional Ni-NTA chromatography.

### Proteolysis assays

*In vitro* proteolysis assays were carried out at 30 °C in 384-well black NBS low-binding plates (Corning), using 0.25 µM ^Eco^ClpX_6_, 0.50 µM ^Eco^ClpP_14_, 2.5 mM ATP, and a regeneration system comprising 16 mM creatine phosphate (MP Biomedicals) and 0.32 mg/mL creatine phosphokinase (Sigma), in PD buffer, in a total reaction volume of 40 µL. Degradation of each GFP substrate was measured by monitoring loss of 511 nm emission following 488 nm excitation in a Tecan Spark plate reader. Initial velocities of enzymatic degradation were fit to a Michaelis-Menten equation in Prism (GraphPad).

### Library construction

Libraries were cloned using saturation mutagenesis [50]. NNK codons were introduced in a forward primer that amplified the region downstream of VapC to avoid introducing inactivating mutations to the toxin at the PCR step. The tag library was cloned into pBAD33 in frame with VapC using BsaI and NcoI endonucleases (NEB) and T4 DNA ligase. For the VapC^YALAX^ library, 40 ng of ligated library plasmid was transformed into MC1061^WT^ by electroporation, and cells were grown in 5 mL 1.5xYT with 25 µg/mL chloramphenicol overnight at 30 °C prior to screening. For VapC^5X^, each MC1061 strain was transformed with ∼1 µg of ligated library plasmid.

The survival rate for each library was determined by measuring the transformation efficiency on selective plates with and without 1% arabinose. Transformants were grown to mid-log phase in 500 mL 1.5xYT with 25 µg/mL chloramphenicol and 0.2% (w:v) glucose at 37 °C. 250 mL of each culture was harvested and maxi-prepped, and the other 250 mL was centrifuged and resuspended in SOB media with 15% glycerol to a concentration of 1×10^10^ cells/mL. Resuspended library transformants were aliquoted into cryovials, flash-frozen in liquid nitrogen, and stored at −80 °C.

### Library screening

VapC^YALAX^ was screened by sub-culturing the overnight library transformants into 5 mL 1.5xYT with and without 1% arabinose for 6 hours at 37 °C. Plasmid from screened and unscreened cells were harvested, purified, and sequenced by Sanger sequencing (Genewiz).

The initial screen of VapC^5X^ in wild type MC1061 cells was carried out in 4x 1L cultures of 1.5xYT with 25 µg/mL chloramphenicol and 1% arabinose, each of which was inoculated with one of the frozen library aliquots described above. A single 1-L uninduced culture was grown. Toxin expression was induced at 37 °C. Plasmid DNA from 2×10^10^ cells was harvested and purified approximately every 2 hours. Maxi-prepped plasmid that was harvested immediately after transformation was used for the 0-hour time point. Purified plasmid was digested with BamHI and HindIII to excise a 545-bp fragment encoding the promoter, VapC and its tag, which was gel-purified prior to Illumina sequencing.

Replicate screening of VapC^5X^ in the proteolytic deletion strains was done as described for the initial screen in wild-type cells. Each library was screened in batch culture and harvested after ∼6 hours of induction or ∼6 doublings (whichever occurred first). Sequence reads were uploaded to the NCBI BioProject repository, ID PRJNA1067636.

### Deep sequencing & analysis

DNA samples from each time point and strain were prepped with a sparQ DNA library prep kit (Quantabio). 2×300-bp paired-end reads were performed on an Illumina MiSeq at the University of Delaware Sequencing and Genotyping Center. A tagseq analysis was performed on the sequencing data by the University of Delaware Center for Bioinformatics and Computational Biology Bioinformatics Core, to exclude reads with mutations to the promotor or toxin open reading frame, or had early stop codons in the tag (**Fig. S2B**). Sequence logos of enriched tags were generated using pLogo [51], scored against the amino acid occurrence within the *E. coli* proteome.

### Growth Assays

Spot-plating assays were carried out by transforming pBAD33 plasmids bearing each VapC variant into MC1061^WT^ or MC1061*^ΔclpAX^*. Transformants were grown overnight at 30 °C in 5 mL 1.5xYT media with 25 µg/mL chloramphenicol (GoldBio) and sub-cultured 1:100 and grown at 37 °C until mid-log phase (∼2 hours). A 10-fold dilution series of cells was made after being diluted to 1×10^7^ cells/mL, 10 µL of which was plated on selective media with and without 1% arabinose (GoldBio) and grown overnight at 37 °C.

Growth rate assays were done by preparing the cells in the same manner described for the spot-plating assays to the point of sub-culturing. Sub-cultured cells were diluted to an OD_600_ of

0.05 in the presence or absence of 1% arabinose. Cell growth was monitored by measuring the OD_600_ of each sample in a 96-well plate over time at 37 °C in a Tecan Spark plate reader.

## RESULTS

### Construction of a selection-based degron discovery platform

We sought to engineer a selection mechanism that couples proteolysis of a target protein to cell growth (**Fig. 1C**). Selection pressure was achieved through expression of the small (∼15 kDa) protein toxin VapC, from the VapBC toxin-antitoxin system of *E. coli* strain W (ATCC 9637) [52]. VapC is a PIN-domain endoribonuclease that cleaves the anti-codon loop of the initiator ^fMet^tRNA [53]. Accumulation of VapC without its cognate VapB antitoxin inhibits translation and arrests cell growth [54].

VapC was cloned into the arabinose-inducible plasmid pBAD33 [45]. Strong induction of untagged VapC with 1% arabinose arrested cell growth on plates (**Fig. 2A**). To determine if growth could be rescued through targeted proteolysis of VapC, we appended the well characterized ssrA degron (AANDENYALAA) from the *E. coli* tmRNA ribosomal rescue system to the VapC C-terminus [24]. SsrA-tagged proteins are recognized and rapidly degraded by the protease ClpXP, and, to a lesser extent, by ClpAP, Lon, and FtsH [24, 25, 27, 55]. Induction of ssrA-tagged VapC (VapC^ssrA^) in wild-type cells resulted in growth at normal levels (**Fig. 2B**). Growth did not occur when VapC^ssrA^ was expressed in a strain lacking elements of the ClpXP and ClpAP proteases (*ΔclpX, ΔclpA*), confirming that wild-type cells escape toxicity through proteolysis of VapC^ssrA^. Direct recognition of the ssrA tag by ClpX is known to require only the downstream portion of its sequence [29, 56]. We confirmed that expression of VapC bearing the terminal 5 residues of the ssrA tag (VapC^YALAA^) permits growth in wild-type cells (**Fig. 2C**). Mutation of the terminal alanine to aspartic acid is known to prevent direct recognition of the ssrA tag by ClpX [29]. As expected, expression of VapC^YALAD^ arrested growth similarly to untagged VapC (**Fig. 2D**). Together, these results demonstrate that degradation of VapC, in accordance with established rules of ssrA degron recognition, relieves VapC-mediated toxicity. Moreover, our findings confirm that this selection strategy successfully couples *E. coli* growth to toxin proteolysis.

**Figure 2.**
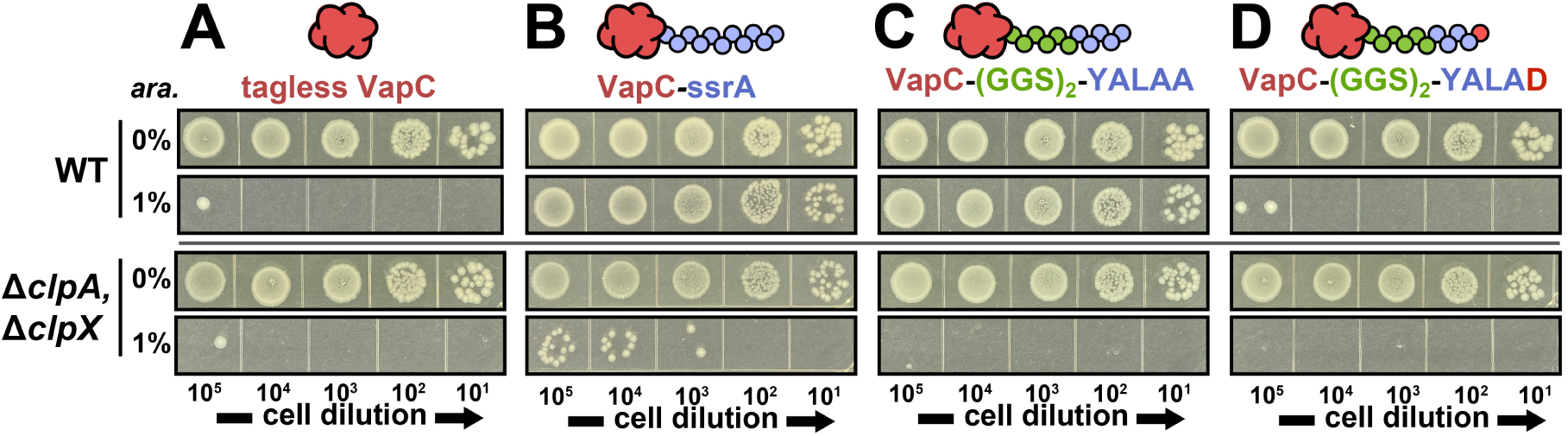
Proteolysis rescues cells from toxicity of VapC expression. **A)** Expression of untagged VapC with 1% arabinose arrests cell growth in wild-type *E. coli* and in a Δ*clpA* Δ*clpX* strain. **B**) Appendage of the full-length ssrA tag to the C-terminus of VapC restores cell growth in wild-type but not Δ*clpA* Δ*clpX* cells. A minimal ssrA tag (YALAA) is sufficient to rescue toxicity in wild-type but not Δ*clpA* Δ*clpX* cells. **D**) No rescue is observed when the terminal Ala is substituted with Asp (YALAD).

### Degron strength correlates with growth rate and fold-enrichment

In order to competitively enrich degrons from a complex library, our screening approach required correlation between degron recognition, cellular levels of VapC, and growth rate. We therefore sought to test whether titration of VapC expression proportionally slows cell growth. We plated cells on media containing 0 – 1% arabinose inducer, and observed that stronger induction of untagged VapC indeed correlates with smaller colony size; colonies were essentially invisible at the highest induction strength (**Fig. S1**). We conclude that VapC levels have a titratable effect on growth rate.

We reasoned that the “strength” of a degron determines the steady-state level of toxin present in the host cell, thereby programming cellular growth rate. In our screen, the rate of VapC proteolysis likely correlates with the *K_M_* with which the degron is processed by cellular proteases. We therefore sought to test the relationship between degron *K_M_* and cellular growth rate. We examined three variants of the minimal ssrA tag: the wild type sequence (YALAA), a conservative A→S substitution in the terminal position (YALA<u>S</u>); and the A→D substitution (YALA<u>D</u>) described above that blocks recognition by ClpX. GFP model substrates bearing these tags were subjected to proteolysis by ClpXP *in vitro*, revealing that ClpXP degrades GFP^YALAA^ with a *K_M_* of 8.3 μM and *k*_cat_ of 1.4 GFP**·**min^-1^**·**enzyme^-1^; GFP^YALAS^ with a *K_M_* of 65 μM and *k*_cat_ of 2.4 GFP**·**min^-1^**·**enzyme^-1^; and GFP^YALAD^ with a *K_M_* of ∼800 and *k*_cat_ of *∼*4 GFP**·**min^-1^**·**enzyme^-1^ (**Fig. 3A**). Thus, increasing severity of mutations to the wild-type tag sequence correlates with increasing *K_M_* for ClpX recognition, but only modest changes in *k*_cat_.

**Figure 3.**
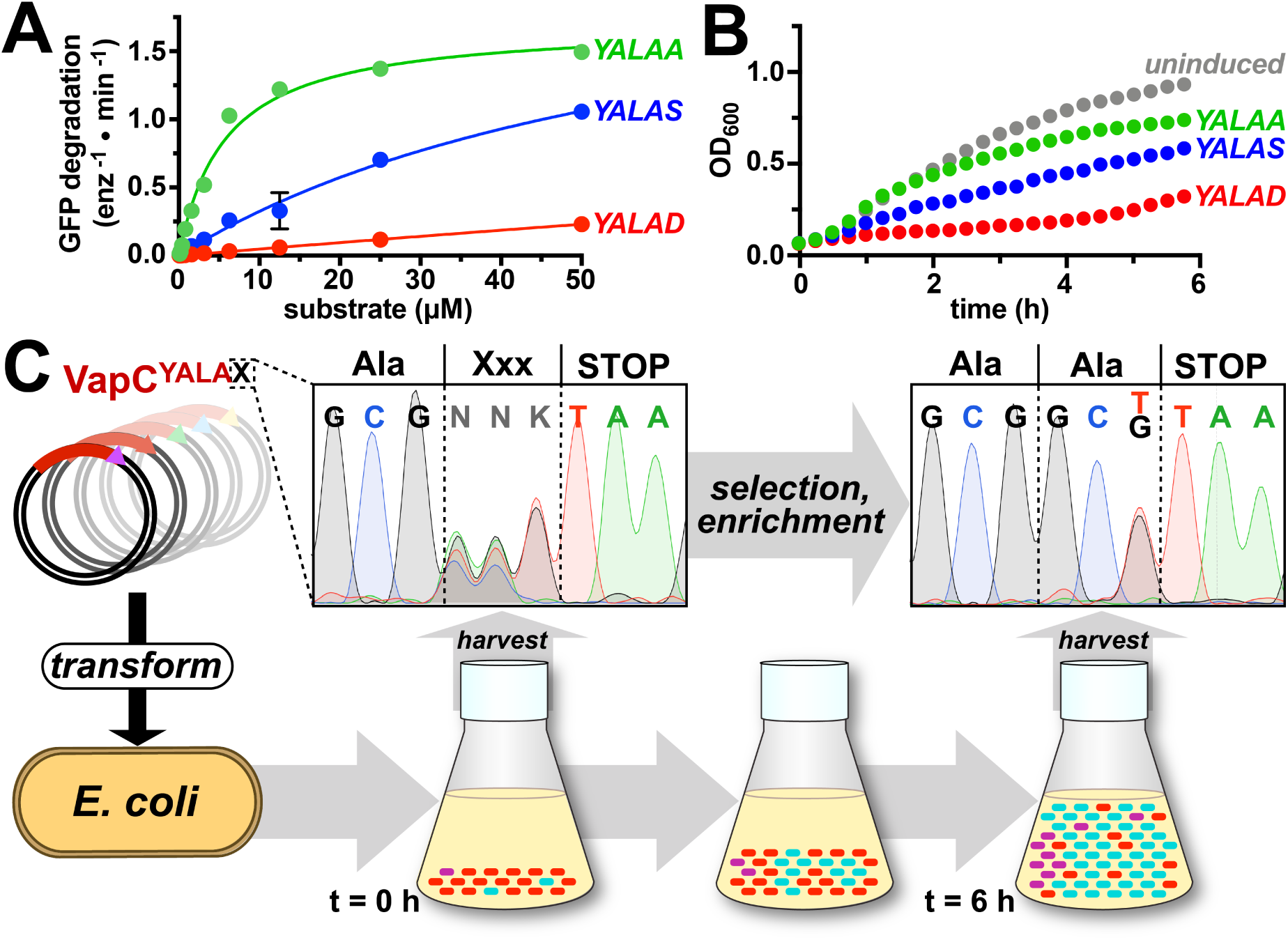
Degron strength correlates with growth rate. **A)** *^Ec^*°ClpXP (0.25 μM) degrades GFP^YALAA^ with a *K_M_* of 8.3 ± 0.9 μM, GFP^YALAS^ with a *K_M_* of 65 ± 18 μM, and GFP^YALAD^ with a *K_M_* of 820 ± 670 μM. Data were fit to a Michaelis-Menten equation. **B**) Expression of VapC^YALAA^, VapC^YALAS^, VapC^YALAD^ in *E. coli* results in growth rates that correlate with each tag’s respective *K_M_*. **C**) A VapC library randomizing the last position of the minimal ssrA tag (VapC^YALAX^) was transformed into wild-type *E. coli* and grown under inducing conditions in liquid culture. Sanger sequencing revealed random codon composition at 0 h but strong enrichment of Ala-encoding codons (GCT, GCG) after 6 h of growth.

We next evaluated how these C-terminal tags influenced the growth rate of cells in the context of VapC expression. By monitoring cell density over time, we found that YALAA supported near wild-type growth, YALAS supported slow growth at ∼50% the wild-type rate, and YALAD supported little growth over the time course of the assay (**Fig. 3B**). These findings suggest that growth rates correlate with degron *K_M_*, and thus with steady-state levels of VapC proteolysis. To confirm that strong degrons are competitively enriched in liquid culture, we created a 20-member library by randomizing the terminal codon in VapC^YALA<u>X</u>^ to NNK (39). This library was transformed into wild-type *E. coli* and harvested after 6 hours of induction with 1% arabinose. Sanger sequencing traces confirmed randomization of the codon at the beginning of the time course, but showed strong enrichment of GGT and GGC Ala codons after 6 hours (**Fig. 3C**). These results demonstrate that the strength of the degron attached to VapC determines tag enrichment in the context of a mixed pool of tags. Thus, by determining fold-enrichment of individual tags in a library, we can rank the relative *K_M_* for proteolysis of individual tag sequences.

### ssrA-like sequences are strongly enriched in wild-type E. coli

We sought to implement the DEtox selection strategy to identify short C-terminal sequences that direct proteolysis of VapC in *E. coli*. To accomplish this, we created a plasmid library encoding VapC, followed by a 6-residue Gly/Ser linker and five C-terminal NNK codons (VapC^5X^), corresponding to a theoretical library size of 3.2×10^6^ unique peptide tags (**Fig. S2A**). The plasmid library was transformed into wild-type *E. coli* MC1061 [57]. Upon plating library cells, we determined that ∼1% of transformants produced colonies under conditions that induce toxin expression (**Fig 4A**). Colonies varied in size, suggesting that individual tag sequences permit different growth rates under VapC^5X^ expression.

**Figure 4.**
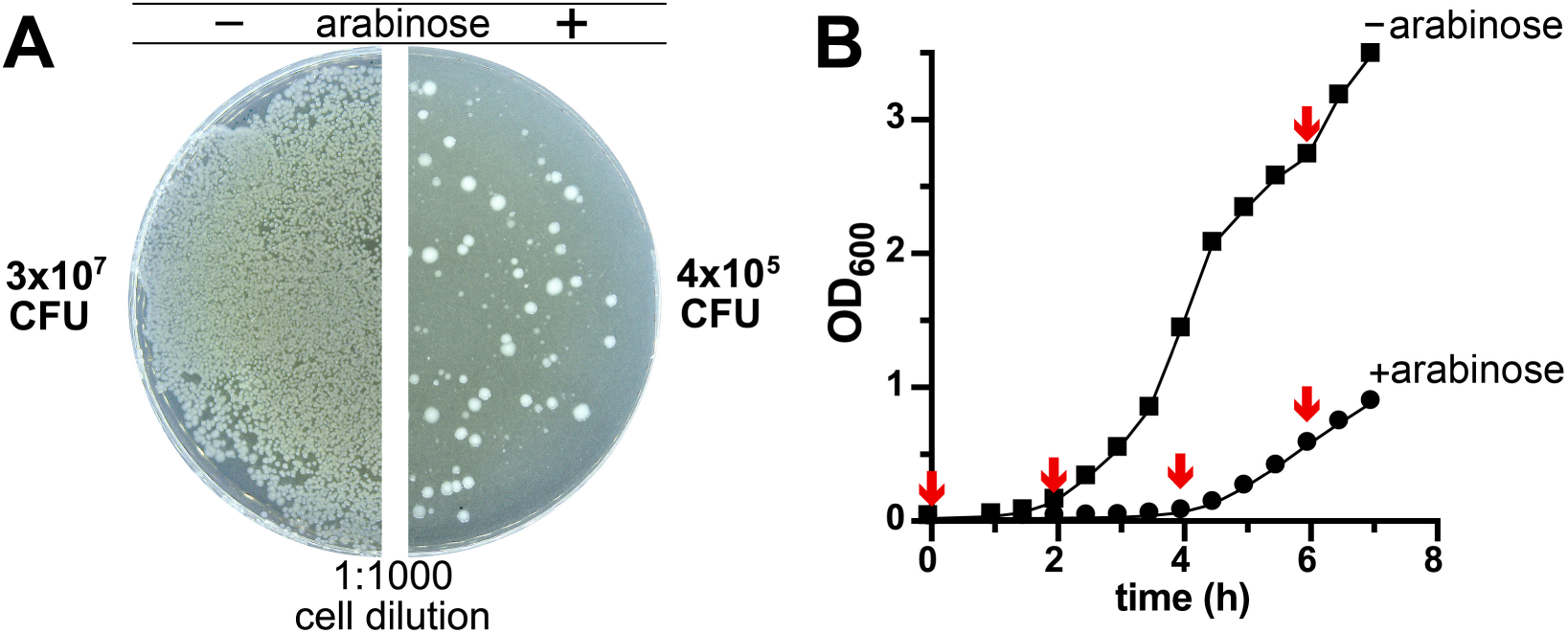
Expression of VapC^5X^ library on plates and in liquid culture. **A)** A VapC library bearing five NNK randomized C-terminal codons (VapC^5X^) was transformed into wild-type *E. coli* and plated on 0% (-) and 1% (+) arabinose. Approximately 1% of library transformants support growth under inducing conditions, producing colonies of varying size. **B**) Cell density of the transformed library was monitored over time in the absence and presence of inducer. Samples of induced culture were harvested at the indicated time points (red arrows) to assess library composition.

The VapC^5X^ library was grown in liquid culture, and toxin expression was induced by 1% arabinose. Cells were harvested prior to induction and at 2, 4, and 6 h post-induction (**Fig. 4B**). As a control, an uninduced sub-culture was grown in parallel and harvested after 6 h. DNA was purified from each sample, and restriction digested to isolate a 545-bp fragment encompassing part of the P*_BAD_* promoter and the entire *vapC^5X^* open reading frame (**Fig. S2A**). Library composition was assessed by paired-end deep sequencing of excised fragments [58]. Sequences with insertions, deletions, or missense mutations were rejected from the analysis (**Fig. S2B**, **Table S1**). (To facilitate meaningful cross-comparisons, sequences with 10 or fewer observations across all samples were pruned from the dataset.) Sequencing captured ∼10 million raw reads across all samples, corresponding to ∼3.1 million reads after filtering, mapped to ∼100,000 unique peptide tags. This represents ∼3% of the total theoretical library. The majority of tags (∼93% of unique sequences) were 5 residues in length, but tags of shorter length were also observed, including all possible 1-mer and 2-mer sequences (**Fig. S2C**). At 6-hours post-induction, about 1000 tags were enriched at least 100-fold, similar to the ∼1% survival rate observed on selective plates (**Fig. 4A, 5A)**. Fold-enrichment correlated well with growth rates estimated from a semi-log plot of tag abundance across time points (**Fig. 5B**). Highly enriched sequences exhibited particularly clear exponential growth, as evidenced by *R^2^* values near 1 (**Fig. S3A**).

**Figure 5.**
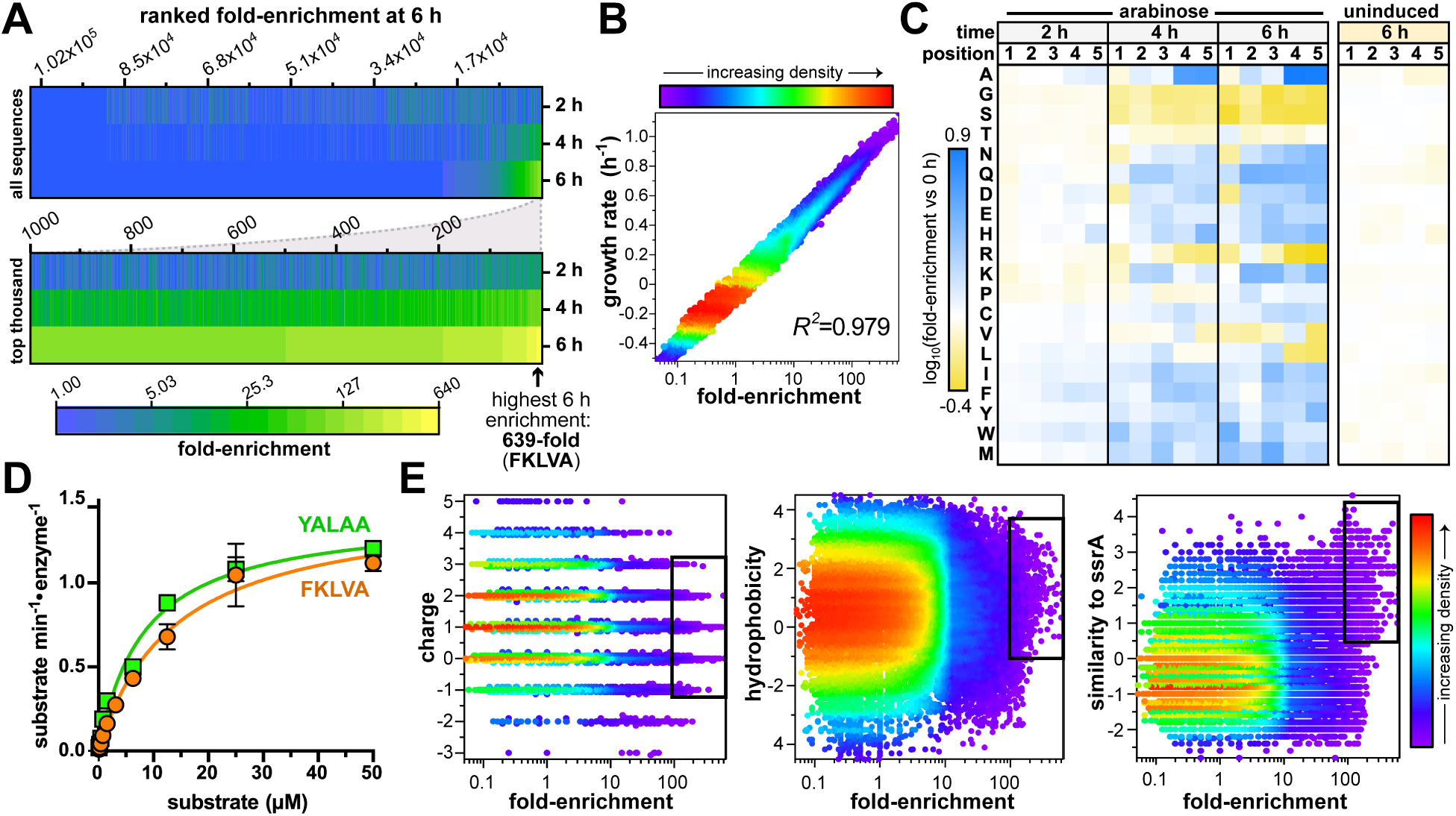
DEtox screening in wild-type *E. coli*. **A)** A heat map illustrates the fold-enrichment of unique sequences compared to their abundance at 0 h. The top ∼1% of sequences were enriched at least 100-fold at 6 h. **B**) The apparent growth rate correlates with fold-enrichment on a semi-log plot. **C**) Heat map of amino acid log fold-enrichment at each tag position over time. (Raw abundance values are reported in **Fig. S3C**). A GFP model substrate bearing the mostly strongly enriched tag, GFP^FKLVA^, is degraded by *^Eco^*ClpXP (0.25 μM) with a similar *K_M_* to GFP^YALAA^ (13.6 ± 1.7 μM and 8.3 ± 0.9 μM, respectively). **E**) Heat maps plot sequence fold-enrichment against net charge, hydrophobicity, or similarity to the ssrA tag (as average per-residue BLOSUM62 score against YALAA). Boxes indicate the most enriched tags, which are comparatively hydrophobic and more similar to the ssrA sequence than the bulk population.

We examined the positional occurrence of amino acids within tags over the course of the experiment. The distribution prior to induction differs from that expected for NNK randomization (**Fig. S3B,C**): several polar residues were comparatively depleted in positions 2 - 5, while Ile, Phe and Tyr were depleted in positions 3 - 5. This trend was consistent across time points, and may reflect compositional biases present in the primers used for cloning. Interestingly, we observed strong shifts in amino acid abundance over the induction time course, which did not occur in the 6-hour uninduced control (**Fig. 5C, S3C**). The most striking change was a ∼7-fold increase in the prevalence of Ala in positions 4 and 5 after 6 h of induction, reaching ∼40% of overall occurrence at these positions. Other amino acids exhibited less pronounced changes: ∼55% depletion of Arg in positions 4 and 5, ∼50% depletion in Gly and Ser in all positions, ∼40% depletion of Leu and Val in the final position, and an increase in abundance of most other amino acids in positions 3 through 5. The lack of Gly and Ser residues may reflect difficulty for unfoldase loops to “grip” these residues, and corresponding slippage during power strokes [59–61].

The ssrA-derived sequence (YALAA) was 118-fold-enriched at 6 h, compared to its initial abundance, confirming that our strategy can successfully enrich a known degron from a complex library. However, the ssrA-derived tag did not appear among the 100 most enriched tags, and instead was ranked 609^th^ (**Fig. S4A,B**). For comparison, the weaker YALAS was enriched only 19-fold (ranked 5465^th^), while the nonfunctional YALAD sequence was not observed at all. The tag with the highest fold-enrichment was FKLVA, enriched 639-fold at 6-hours (**Fig. 5A**). While this tag *per se* has not been reported previously to our knowledge, a terminal LVA sequence was observed in a prior screen for C-terminal tags that destabilize GFP in *E. coli* [62], and is known to be recognized by ClpX [63, 64]. Indeed, a GFP^FKLVA^ model substrate was degraded *in vitro* by *^Eco^*ClpXP with *K_M_* and *k*_cat_ similar to GFP^YALAA^ (**Fig. 5D**).

### Upstream composition influences recognition of ssrA-like tags

Strongly enriched sequences tended to be neutral or carry a positive charge ≤2, were moderately hydrophobic, and bore clear similarity to the ssrA-derived tag (**Fig. 5E**). In particular, terminal Ala-Ala motifs occurred in 51% of the top 1000 sequences and 91% of the top 100 sequences (**Fig. 6A**), in accordance with the enrichment of Ala at positions 4 and 5 in the bulk analysis (**Fig. 5C; Fig S3C**). However, terminal Ala-Ala motifs alone were not sufficient to drive strong enrichment. 39% of sequences with a terminal Ala-Ala were enriched less than 19-fold (below the level of YALAS), and 20% reached 5-fold enrichment or below.

**Figure 6.**
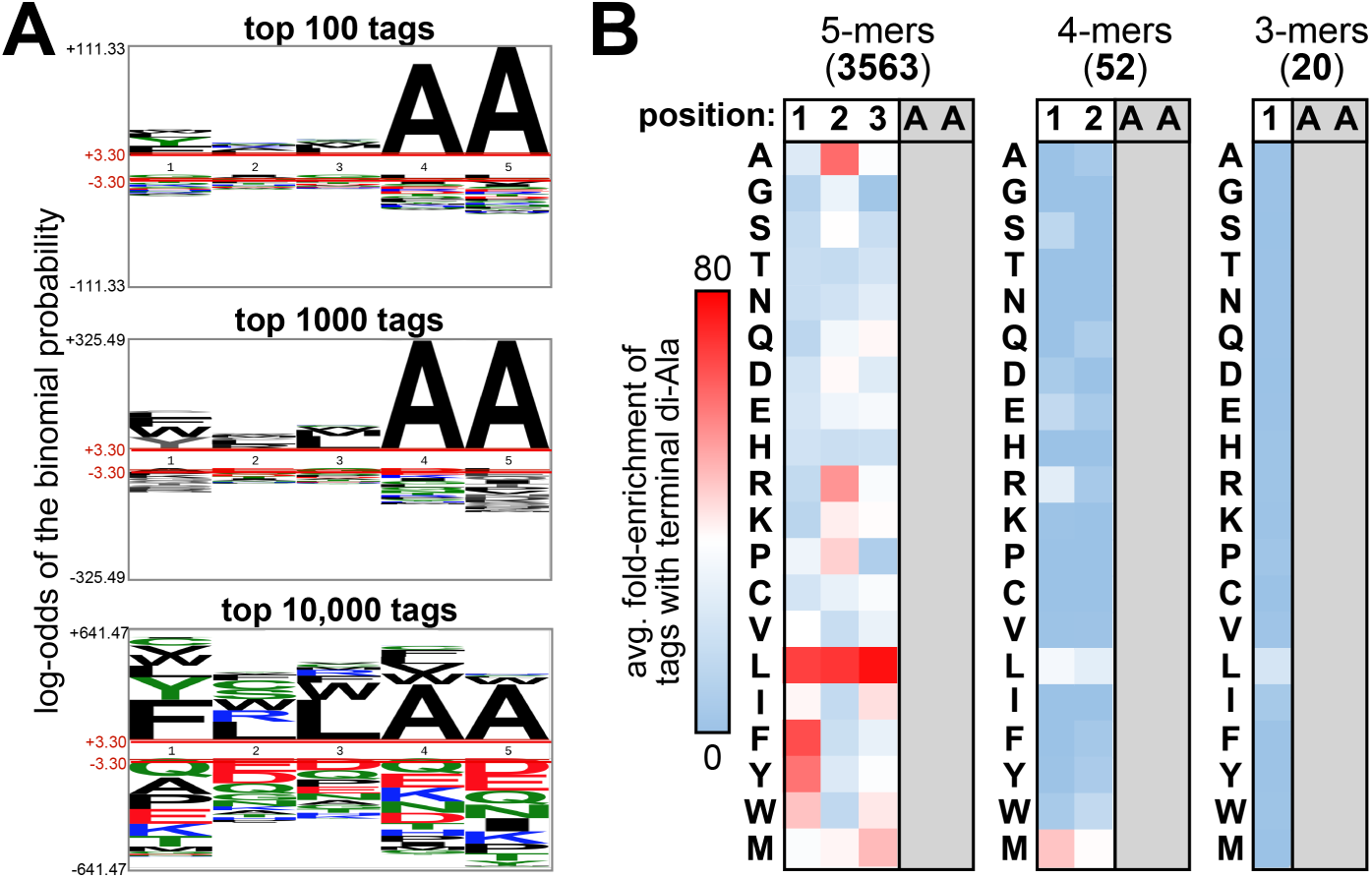
Terminal Ala-Ala motifs are abundant among enriched 5-mer tags. **A)** Sequence logos of the full library, the top 1000, and the top 100 highest fold-enrichment tags. **B**) Heat map illustrating the average fold enrichment of 5-mer, 4-mer or 3-mer tags ending in Ala-Ala, bearing the indicated amino acid at the indicated position.

To understand how upstream residues contribute to recognition of tags ending in Ala-Ala, we correlated upstream amino acid occurrence with average fold-enrichment (**Fig. 6B**). Among 5- mer sequences there was clear compositional bias. Leu in upstream positions strongly favored enrichment, as did Ala or Arg at position 2, and aromatic amino acids in position 1, resulting in a consensus motif of [Leu/Phe/Tyr/Trp]-[Leu/Ala/Arg]-Leu-Ala-Ala. Conversely, some residues were depleted, including polar amino acids at position 1; most β-branched and bulky amino acids at position 2; and Pro, His, Gly and amino acids with short polar sidechains at position 3. Interestingly, gammaproteobacterial ssrA tags overall have more restricted sequence variation than this consensus motif (**Fig. S3D**), with “YALAA” by far the most common [65]. Evolutionary conservation of tmRNA elements may thus be driven by physiological interactions and constraints that are not recapitulated in our screen.

Our dataset also included shorter tags that terminate in Ala-Ala. In contrast to 5-mers, no 4-mer, 3-mer or 2-mer terminating in Ala-Ala experienced strong enrichment, and there was only weak correlation between upstream amino acids and enrichment level (**Fig. 6B**). The presence of an upstream Leu, for example, had only a small correlation with enrichment among 4-mers and 3- mers. (The strongest positive association was observed with Met in position 1 of 4-mers.) Notably, a dimer of Ala-Ala alone was entirely defective, leading to depletion rather than enrichment. The loss in function in shorter tags is likely due in part to the Gly/Ser linker, which positions GGSGGS upstream of the tag. Indeed, all observed 5-mer sequences with Gly/Ser preceding Ala-Ala (SSSAA, SSGAA, GSSAA, GSGAA, and SGSAA) had low fold-enrichment, ranging from 0 – 2- fold at 6 h. Thus, both tag length and upstream amino acid composition influence functionality, even among those ending in Ala-Ala.

Several highly enriched sequences in the dataset were starkly dissimilar to the ssrA tag, and we re-cloned and individually tested these for the ability to rescue VapC toxicity on plates. None of the non-ssrA-like tags rescued growth under inducing conditions (**Fig. S4**), suggesting that their enrichment was driven by mutations outside of the sequenced *vapC* cassette. This reinforces the observation that the majority of strongly enriched sequences were ssrA-like, and that few degrons of similar strength deviate from this template. The appearance of strongly enriching false-positive hits is likely a consequence of the strong selective pressure favoring background mutations that nullify VapC toxicity.

### Screening in protease deletion strains

The results of the DEtox screen highlight several limitations to degron screening in wild-type *E. coli*. The preponderance of strong ssrA-like tags, which are likely ClpXP-dependent, complicates identification of less potent tags. Additionally, the presence of the full complement of endogenous proteases introduces ambiguity over which proteases recognize particular degrons. We sought to address both limitations by performing parallel screens in wild-type cells and in deletion strains lacking components of ClpXP or ClpAP (a deletion of the entire *clpS-clpA* operon, *ΔclpSA*; *ΔclpX*; and *ΔclpP*) which carry out a large portion of protein turnover [55, 66].

The VapC^5X^ library produced proportionally fewer transformants in strains harboring deletions of proteolytic components, compared to wild-type (**Fig S5A**). The fewest viable transformants occurred in *ΔclpX* and *ΔclpP* cells, suggesting that most functional degrons in wild-type cells are recognized by ClpXP. Growth rates for each strain in liquid culture correlated with their respective survival rates on plates (**Fig. S5B**).

Screens of VapC^5X^ in wild-type and mutant strains were carried out in two biological replicates, and library composition was assessed at 0 and 6 h post-induction by multiplexed paired-end deep sequencing (**Table S2**). Multiplexing reduced the number of reads, with each sample capturing ∼0.2% of the theoretical library. To filter out false positives arising through mutations beyond of the *vapC* locus, we focused on hits enriched by at least 5-fold in both replicates (**Fig 7A**). Only a minority of sequences passed these criteria. For example, 2,155 and 1,522 sequences were enriched by at least 5-fold in the individual wild-type replicates, but only 611 sequences met the enrichment cutoff in both. Fewer consistently enriched tags appeared in strains harboring protease disruptions compared to wild-type, in line with the pattern of growth observed on plates and in bulk culture under inducing conditions (**Fig. S5**).

**Figure 7.**
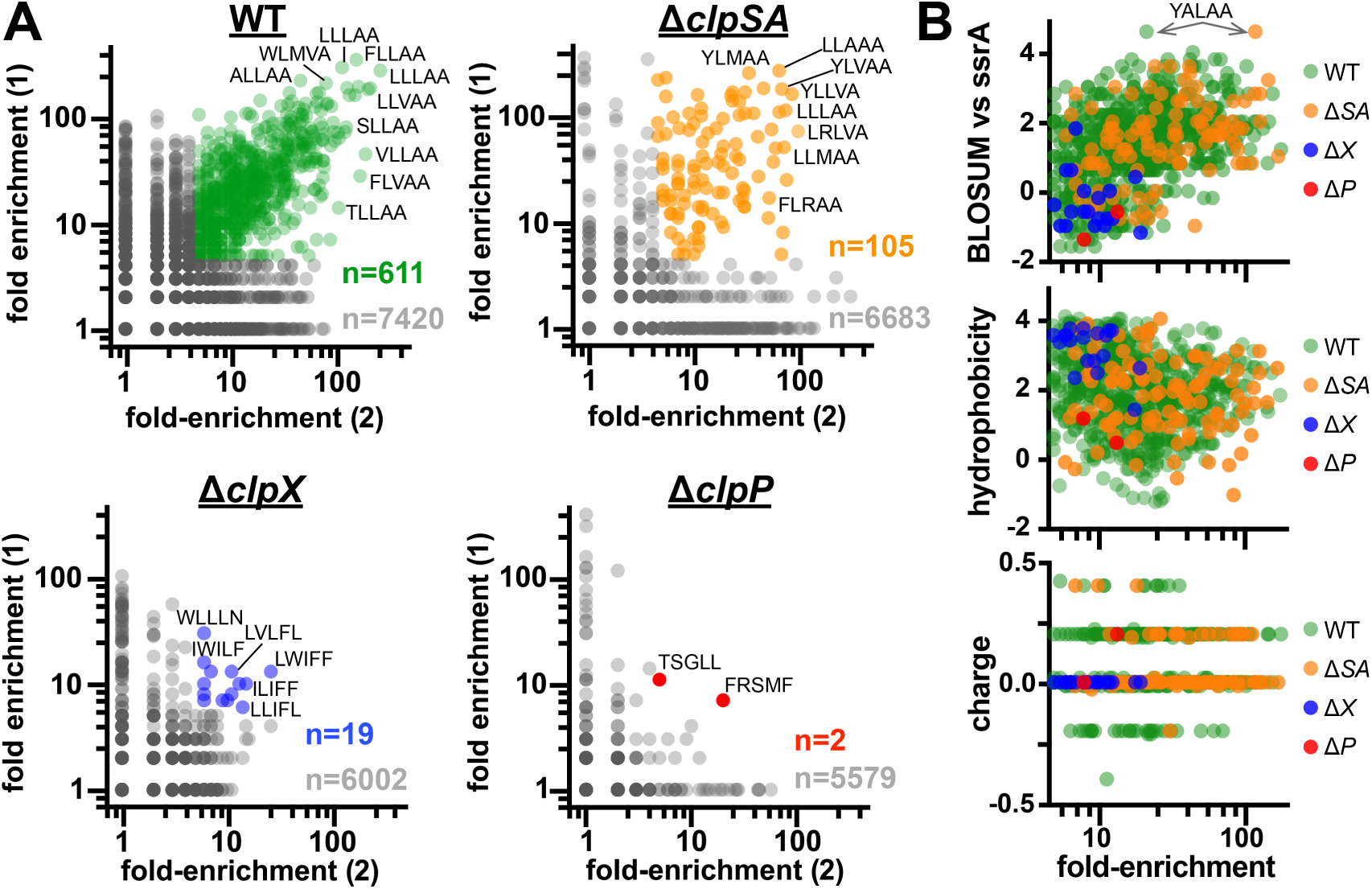
Fewer strongly enriched degrons appear in strains lacking components of ClpXP. **A)** Fold-enrichment of tags observed across replicates in the indicated *E. coli* strain were cross-compared to identify consistently enriched tags. Tags enriched after 6 hours but not observed in the pre-induction samples were set to an initial occurrence of 1. Tags enriched <5-fold in at least one replicate are colored gray, and make up the majority of observed sequences in each strain. **B**) Charge, hydrophobicity and ssrA similarity of consistently enriched tags are plotted against fold-enrichment.

Sequences that were consistently enriched in strains with an intact ClpXP protease – wild-type and *ΔclpSA* – were predominantly ssrA-like, somewhat hydrophobic, and neutral to slightly positively charged (**Fig. 7B**). Notably, fewer enriched tags were observed in *ΔclpSA* than in wild-type, although they bore similar ssrA-like character. Similarly, the ssrA-like “YALAA” tag was enriched to a lower level *ΔclpSA*, perhaps due to the slower growth rate of this deletion strain (**Fig. S5B**). Few enriched sequences emerged in *ΔclpX* and *ΔclpP* strains, both of which lack intact ClpXP. Sequences that did enrich in these strains reached lower fold-enrichment and lacked obvious similarity to ssrA. Non-ssrA-like tags that consistently enriched in *ΔclpX* were scrutinized individually. These bulky, hydrophobic tags (LWIFF, LVLFL, IWILF, & LLIFF) supported intermediate growth as VapC fusions in all proteolytic deletion strains tested (**Fig. S6A**). However, the tags did not promote proteolysis of a GFP model substrate *in vitro* by ClpXP or ClpAP (**Fig. S6B**). It is possible that these sequences compromise VapC folding and solubility, or mimic inhibitory interactions made by hydrophobic segments of the VapB antitoxin that block VapC activity [52, 67].

## DISCUSSION

ATP-dependent proteases recognize some substrates via degrons, but the prevalence of simple degron sequences and their corresponding proteolytic strength has remained an open question. Most known degrons were identified through genetic or capture-based approaches that are fundamentally restricted to the limited sequence space of the proteome. Through the use of randomized terminal sequences, the cell-based DEtox screening method implemented here allowed us to explore a substantially larger segment of sequence space, and develop a more expansive picture of the relationship between C-terminal sequences and proteolysis in *E. coli*.

Remarkably, we only found consistently enriched 5-mer tags that restored near-normal growth in strains possessing ClpXP. These ClpXP-dependent degrons bore similarity to the ssrA tag, with a strongly conserved terminal Ala-Ala motif. Although *E. coli* possesses five ATP-dependent proteases, four of which are capable of recognizing the ssrA tag [16, 23–27], our data suggest that the interaction between ClpXP and ssrA-like sequences is a singularly robust recognition modality among short C-terminal degrons. The preferential enrichment of ssrA-like tags likely reflects the importance of ribosome rescue in cellular fitness. Ribosome stalling occurs regularly and must be resolved quickly and efficiently [16, 68, 69]. The affinity of ClpX for ssrA-like tags provides an evolutionarily conserved mechanism to prioritize proteolysis of incomplete polypeptides liberated from stalled ribosomes.

The preponderance of ssrA-like tags in our dataset allowed us to assess the positional influence of amino acids on tag function. As expected from prior studies [29, 56], the terminal Ala-Ala motif is critical. Upstream positions also contribute to functionality, although the pattern of amino acid preference differs from the conserved “YALAA” sequence found among gammaproteobacterial ssrA tags. For example, Leu in any upstream position correlated with enrichment in our screen, whereas Leu occurs only in the antepenultimate position of ssrA. More surprisingly, Arg in the 2^nd^ position was also associated with proteolysis, yet Arg is virtually absent from proteobacterial ssrA sequences. These differences suggest that the full physiological ssrA tag evolved to satisfy more complex constraints than imposed by our selection scheme. Indeed, upstream elements of ssrA are important for recognition by ClpA and the ClpX adapter SspB [29, 70, 71]; other components may be important for recognition by Lon or FtsH. Recognition of ssrA by diverse proteases may aid degradation of polypeptides that are difficult for ClpXP to process. By contrast, our selection scheme prioritized rapid and efficient proteolysis, which may be best accomplished by ClpXP. The discrepancy between effective degron composition and sequence conservation highlights the utility of direct screening for predicting proteolytic susceptibility, and for guiding degron engineering in synthetic biology applications [33, 34].

Our dataset also reveals features within ssrA-like tags that impede degradation. Gly and Ser in upstream positions prevent VapC proteolysis, which may reflect a defect in unfolding or recognition. Unfolding by ClpX is impaired by Gly-rich sequences in positions important for grip and pulling [59–61]. However, studies with GFP demonstrate that residues 3 – 5 positions away from the folded protein are most critical for application of force-generating power strokes [60], whereas the upstream positions in our tag library are at least 7 – 9 residues from VapC, suggesting that impaired proteolysis here is not related to unfolding. Rather, it is likely that upstream Gly and Ser weaken initial tag recognition or deprive ClpX of a tractable site for application of the initial power stroke. Our findings also confirm prior data on the minimal length of a functional ssrA-like tag [28]. The terminal di-Ala motif is necessary for recognition, but not sufficient: no tag shorter than 5 residues was robustly enriched in our dataset, suggesting that positioning of GGSGGS nearer the terminal Ala-Ala abrogates tag engagement.

Perhaps the most surprising finding in our dataset was the stark absence of strongly enriching non-ssrA-like and ClpXP-independent tags. To some extent, this may reflect biases in the screening approach, which was benchmarked against VapC^ssrA^ and thus optimized to identify degrons of similar strength. The live-or-dead nature of the screen necessitated high levels of VapC expression, which may overwhelm some proteases. Enrichment of ClpXP-independent tags may require supplemental expression of individual proteases in the context of *clpX* deletion. Nevertheless, our overall results have implications on the shape of the proteolytic landscape in *E. coli* and other bacteria. If short non-ssrA-like degrons exist, we infer that their recognition has been calibrated by evolution to be weak, favoring protein stability and allowing evolutionary variation of terminal sequences without frequent intrusion on degron sequence space. This is consistent with findings that protein C-termini across bacterial taxa are biased away from hydrophobic residues correlated with proteolytic susceptibility [42]. Efficient proteolysis of proteins lacking ssrA-like tags likely requires more complex modes of recognition, such as longer degron sequences, multivalent recognition elements, or delivery by proteolytic adaptors. Such complex recognition paradigms may be inherently advantageous by providing more opportunities for regulation [2, 10], thereby allowing cells to tune proteolytic programs to a wider range of conditions [8].

Our findings likely hold true at least among gammaproteobacteria, given the general conservation of ATP-dependent proteases within this clade. Moreover, the selection-based DEtox screening approach can likely be implemented in a variety of bacteria of interest and optimized to interrogate degrons of varying strength and complexity. A comprehensive understanding of degron sequence space may ultimately facilitate the identification of proteolytic pathways in bacterial pathogens, and enable the creation of robust proteolytic circuits in engineered contexts.

## DATA AVAILABILITY

Sequencing datasets were uploaded to the NCBI BioProject repository with ID PRJNA1067636.

## AUTHOR CONTRIBUTIONS

PCB carried out experiments. PCB and KRS conceived of the project, analyzed data, and prepared the manuscript.

## ACKNOWLEDGEMENTS

We thank Cady Burnside and Henry Anderson for assistance with cloning, and all members of the Schmitz lab for helpful comments and advice. We thank Bruce Kingham and Mark Shaw at the University of Delaware DNA Sequencing & Genotyping Center and Shawn Polson at the University of Delaware CBCB Bioinformatics Core for assistance with experimental design, sample preparation, DNA sequencing, and data analysis.

