## Supplementary Figures S1 - S6 for "Toxin-based screening of C-terminal tags in *Escherichia coli* reveals the exceptional potency of ssrA-like degrons"

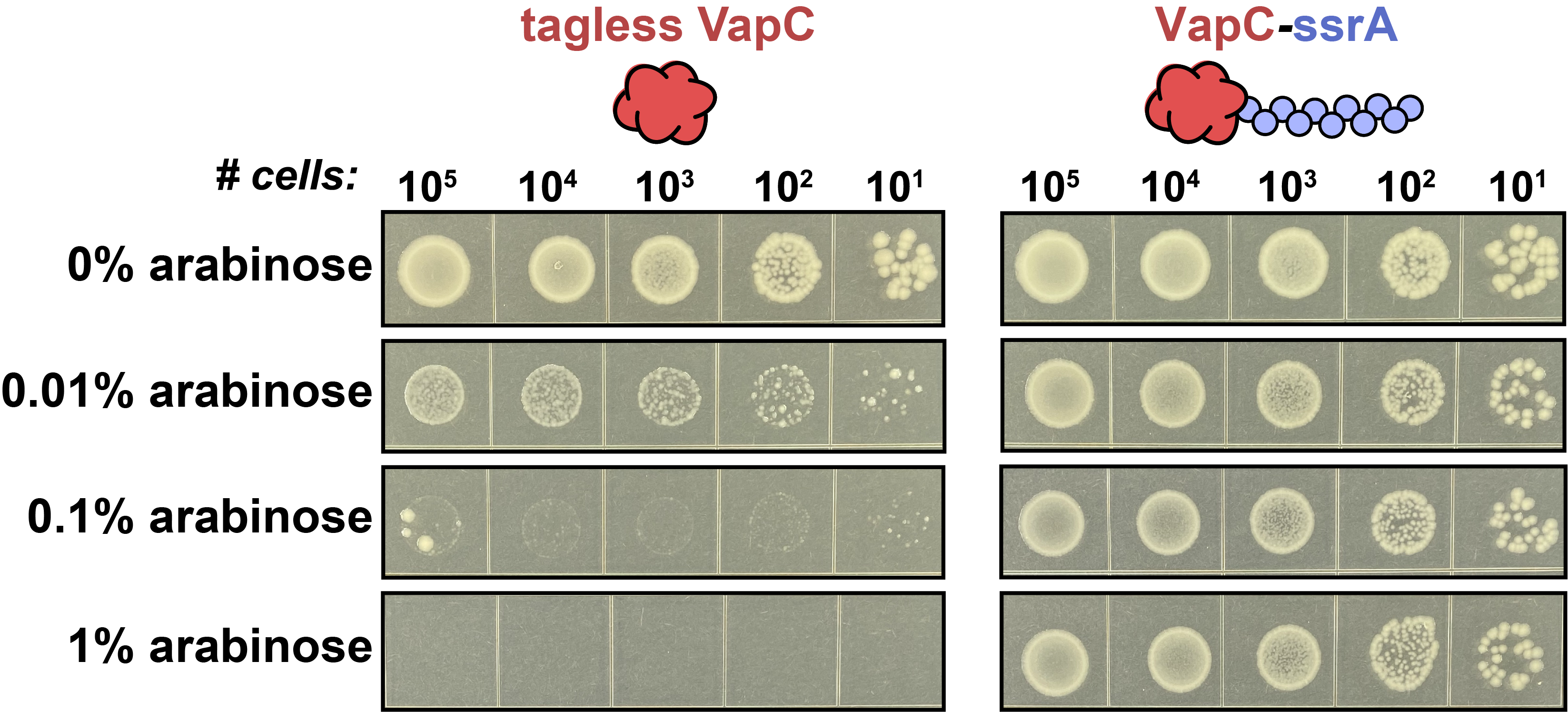

**Supplemental Figure S1. Validation of VapC toxicity and proteolysis.** Untagged and ssrA-tagged VapC were expressed in wild-type *E. coli* on plates with increasing arabinose concentrations.

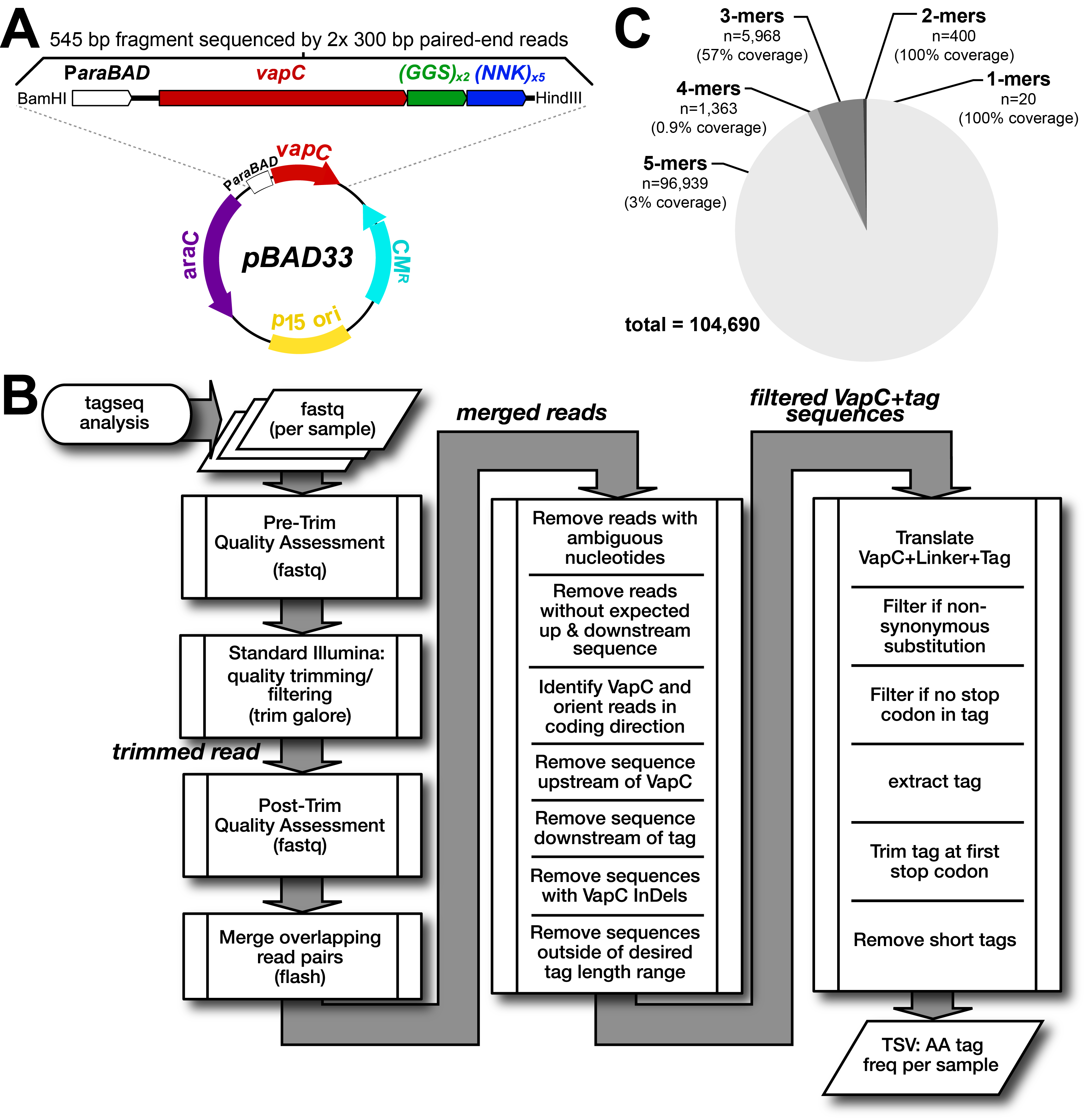

**Supplemental Figure S2. Bioinformatic pipeline for analyzing deep sequencing of plasmid library. A**) Vector map of VapC-pBAD33 showing the region that was sequencing – a 545-bp fragment extending from the end of the P*araBAD* promoter to the stop codon after the tag. This fragment was excised after restriction digestion prior to Illumina sequencing. **B**) Raw Illumina sequencing data was filtered through a bioinformatic pipeline before analysis. **C**) Out of the 104,690 tags identified in our first screen, 93% were 5-mers, 1.3% were 4-mers, 5.7% were 3-mers, 0.38% were 2-mers, and 0.02% were 1-mers. The observed coverage of tags in each category is noted.

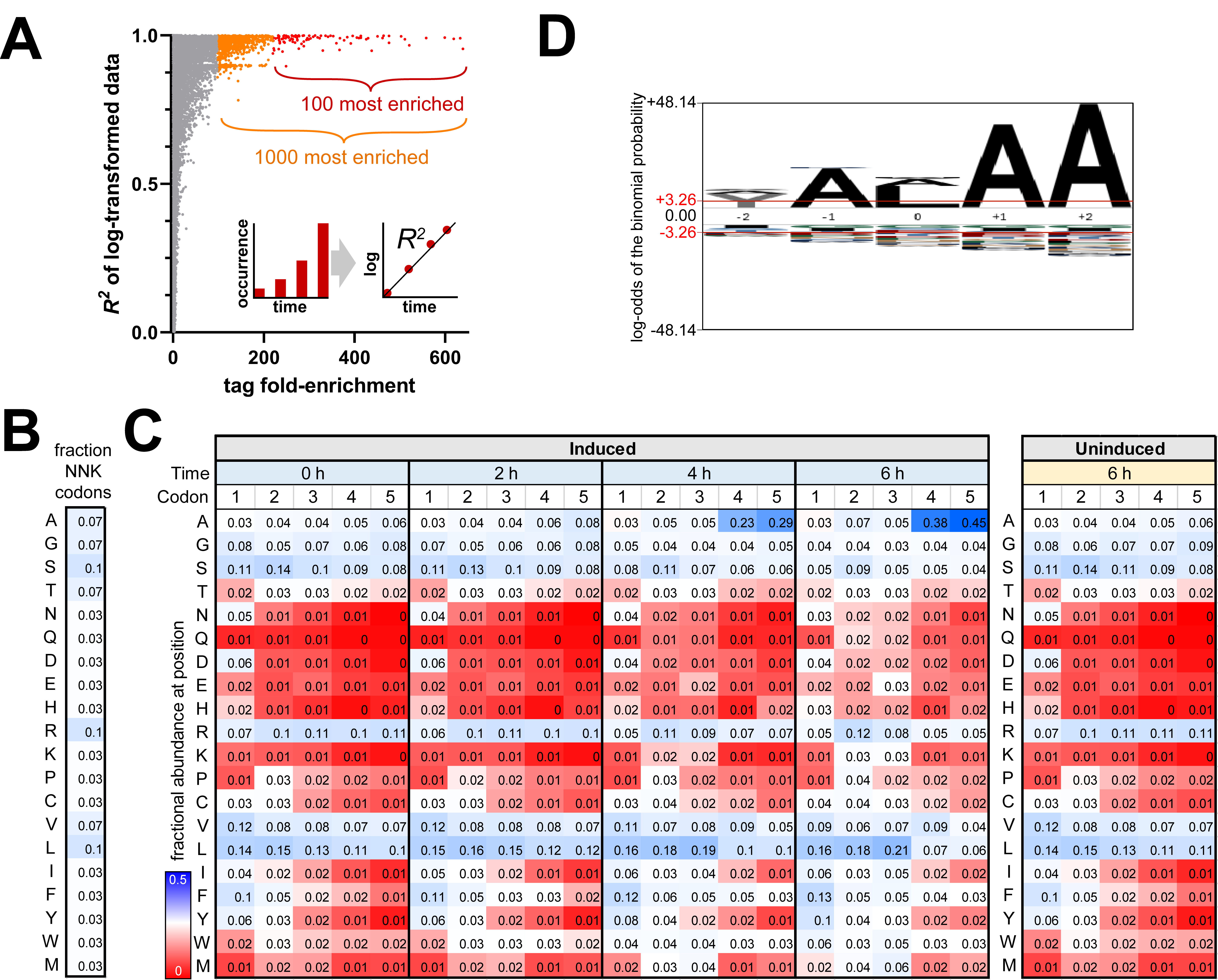

**Supplemental Figure S3. Positional analysis and bulk statistics. A)** Comparison of tag fold-enrichment to the *R^2^* value of linear regression fit to log-abundance values vs time – a metric for assessing consistent growth rate. Highly enriched sequences have *R^2^* values nearer to 1, indicating consistent growth and enrichment over the course of the DEtox experiment. **B**) Heat maps showing the theoretical fractional abundance of amino codons resulting from NNK randomization, and **C**) positional abundance of amino acids over time. **D**) Sequence logo of the last five positions of ssrA tags from 280 gammaproteobacterial species.

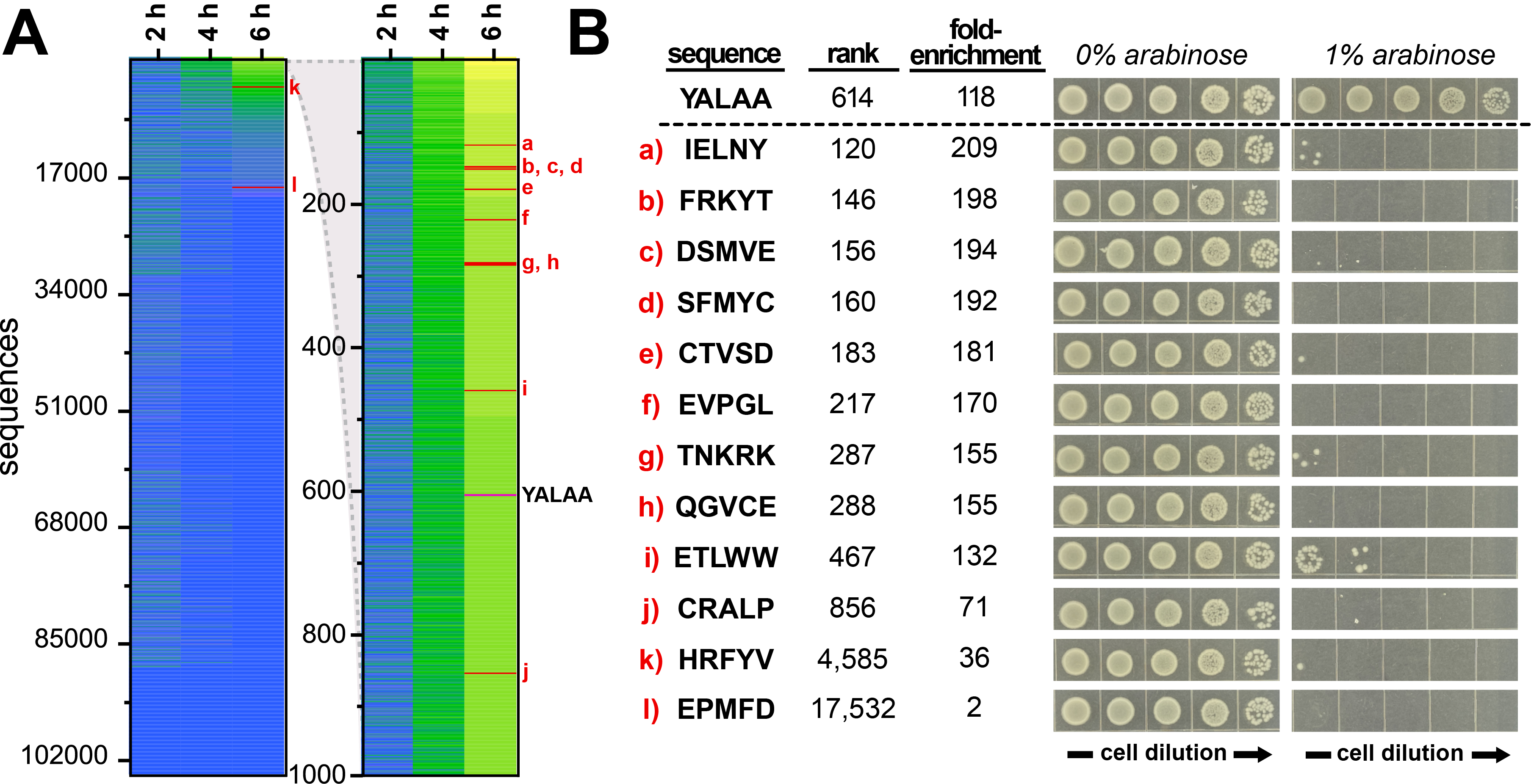

**Supplemental Figure S4. Highly enriched non-ssrA-like degrons are likely false positives*.* A**) A set of non-ssrA like degrons of varying fold-enrichment were selected from the dataset. Their position in the heatmap is indicated, as is the position of the YALAA tag. **B**) Tags were individually cloned onto the C-terminus of VapC, and growth was examined in the absence and presence of inducer. Non-ssrA-like tags were unable to rescue growth in the presence of arabinose.

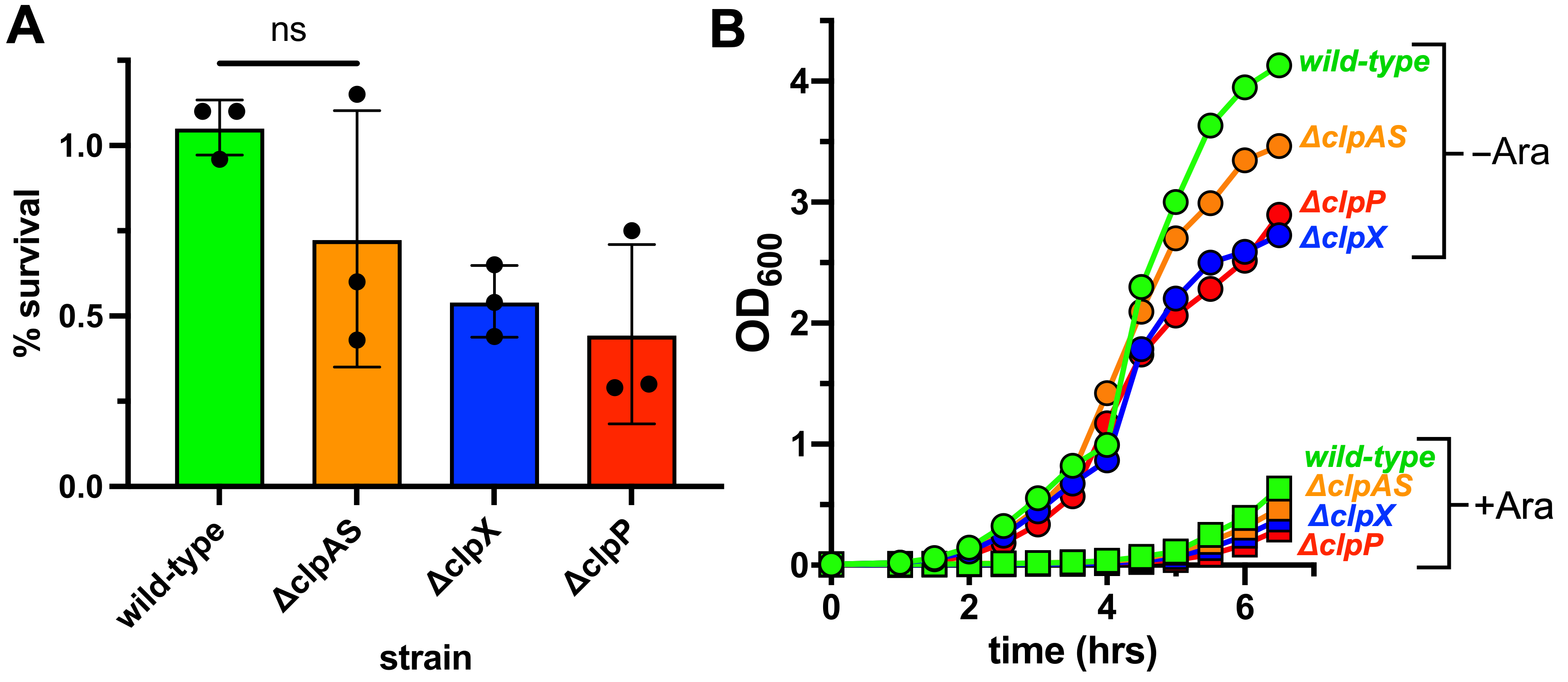

**Supplemental Figure S5. Deletion of protease components reduces survival and growth of VapC^5X^ library transformants. A**) A plasmid library encoding VapC^5X^ was transformed into the indicated *E. coli* strains and colony forming units were measured on plates with and without 1% arabinose. The number of colonies that appeared under inducing conditions is plotted as a percentage of the total colonies observed under non-inducing conditions. **B**) The indicated strains harboring the VapC^5X^ library were grown in liquid culture in the presence and absence of 1% arabinose. Growth was monitored over time by OD_600_.

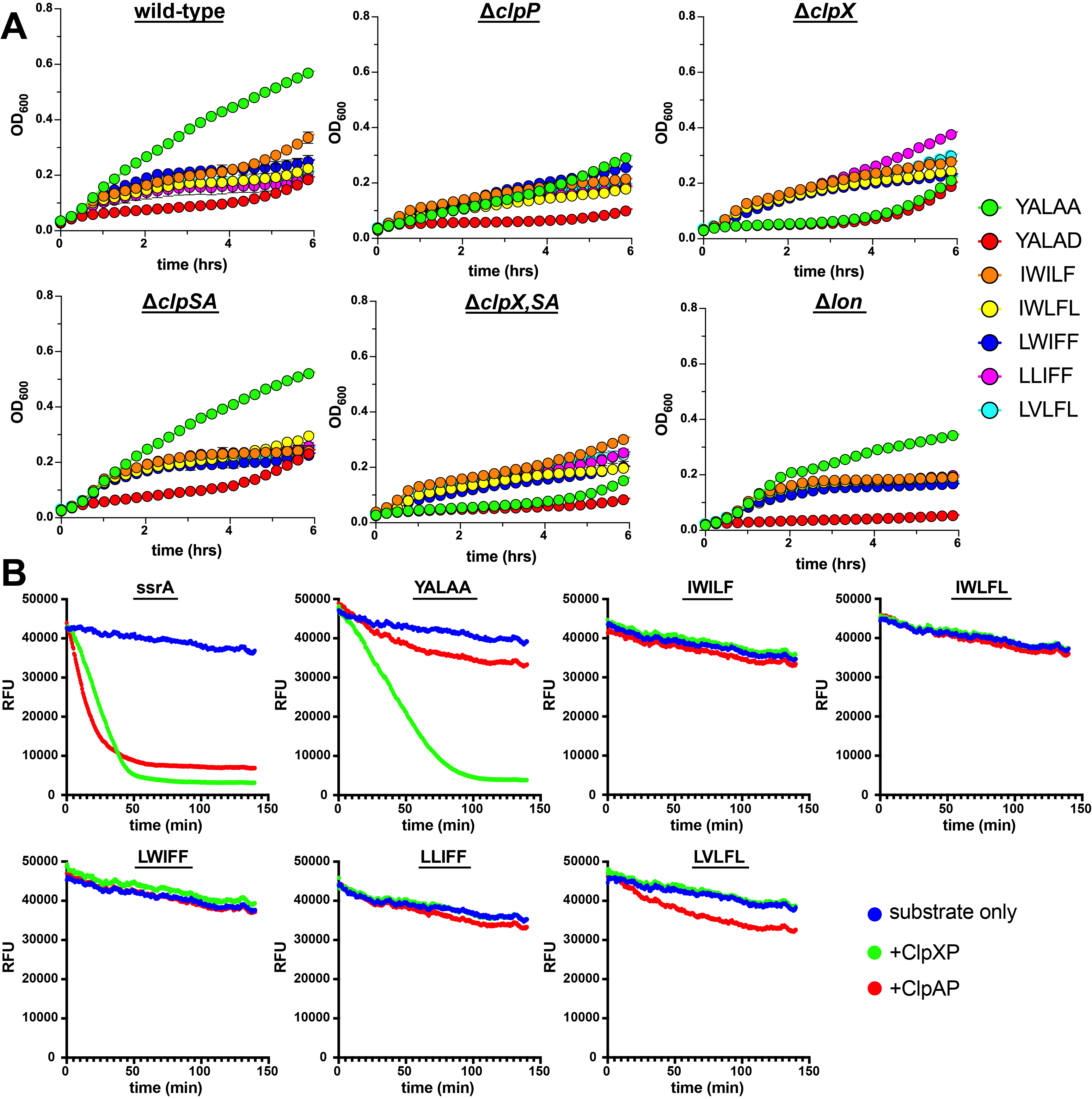

**Supplemental Figure S6. ClpX-independent degrons support slow growth *in vivo* but not proteolysis *in vitro.* A**) YALAA supports robust growth in all strains in which ClpXP is present. YALAD does not support robust growth in any strain. Degrons that were enriched in *ΔclpX* cells (see **Fig. 7**) support at least partial growth in all deletion strains. **B**) Model GFP substrates were constructed carrying the indicated C-terminal tag, and proteolysis of 15 μM model substrate by ClpXP or ClpAP (0.25 μM) was assessed *in vitro*. GFP bearing the full ssrA tag was degraded by ClpXP and ClpAP. GFP bearing the minimal ssrA sequence, YALAA, which lacks sequence determinants for ClpA recognition, was degraded by ClpXP but only modestly by ClpAP. None of the ClpX-independent degrons was degraded by ClpXP, and only GFP^LVLFL^ was modestly degraded by ClpAP.
